# Draft Genome Analysis of *Streptomyces sp*. IICT-RSP475: Unveiling Novel Biosynthetic Potential

**DOI:** 10.1101/2025.03.20.643947

**Authors:** Uma Rajeswari Batchu, Mahesh Anumalla, Chandrasekhar Cheemalamarri, Joshna Rani Surapaneni, Prakasham Reddy Shetty

**Author notes:** Corresponding author & Reprints Prakasham Reddy Shetty. These authors contributed equally (Uma Rajeswari Batchu, Mahesh Anumalla). **Author’s Email ID** Uma Rajeswari Batchu -, Mahesh Anumalla -, Chandrasekhar Cheemalamarri -, Joshna Rani Surapaneni -, Prakasham Reddy Shetty.

## Abstract

*Streptomyces* species are ubiquitous bacteria renowned for their prolific production of pharmaceuticals and therapeutic agents. In this study, we explored the draft genome of a novel *Streptomyces* species, *Streptomyces sp*. IICT-RSP475, isolated from Talakona, Tirupati, using next-generation sequencing and bioinformatics tools. The draft genome of *Streptomyces* sp. IICT-RSP475 consists of 4,550,481 base pairs (bp) with a high GC content of >70%. Genome analysis identified 18 biosynthetic gene clusters (BGCs) responsible for the production of ribosomally synthesized and post-translationally modified peptides (RiPPs), polyketide synthases (PKS), non-ribosomal peptide synthetases (NRPS), and other secondary metabolites, including hydrogen cyanide, terpenes, and N1-siderophores. Notably, genome mining using the antiSMASH tool uncovered two novel BGCs classified as RiPP-like clusters. These clusters exhibited unique genetic architectures with previously uncharacterized biosynthetic genes, suggesting the presence of novel bioactive metabolites. Ribotyping analysis, further supported by TYGS (Type Strain Genome Server) and ribosomal MLST (Multilocus Sequence Typing), confirmed the classification of this strain as a novel *Streptomyces* species. These findings highlight the genomic potential of *Streptomyces sp*. IICT-RSP475 and warrant further investigation into the expression, structural elucidation, and functional characterization of its novel therapeutic metabolites.

## Introduction

*Streptomyces* are widely distributed Gram-positive bacteria in soil, belonging to the phylum Actinobacteria. They possess distinctive characteristics, including an exceptionally high GC content (>70%), large linear genomes (8–10 Mb), and extensive biosynthetic capabilities (Hopwood. 2019). *Streptomyces* species are regarded as natural biofactories, harboring multiple biosynthetic gene clusters (BGCs) responsible for the production of bioactive compounds with significant pharmacological potential (Ward & Allenby, 2018). These bacteria are prolific producers of over 100,000 antibiotics, contributing to 70 to 80% of all natural bioactive products (Abbasi et al. 2020). Beyond antibiotic production, they also synthesize compounds with therapeutic applications in treating life-threatening diseases, including cancer and Alzheimer’s disease. Genome mining is an in silico approach used to identify BGCs encoding novel bioactive compounds with pharmaceutical and agrochemical applications (Zhao et al. 2019). It has been reported that each *Streptomyces* genome contains approximately 25 to 70 BGCs, with diverse biological functions (Challis. 2014). This approach enables the exploration of untapped genomic potential in uncultured *Streptomyces* and facilitates the characterization of novel BGCs in cultured strains through comparative analyses using reference BGCs from the antiSMASH (https://antismash-db.secondarymetabolites.org/) and MIBiG (https://mibig.secondarymetabolites.org) databases (Niu. 2018). To date, antiSMASH has identified 1,346 BGCs across 39 *Streptomyces* genomes, underscoring the biosynthetic richness of this genus. In the present study, we report the isolation of a novel *Streptomyces* strain, *Streptomyces* sp. IICT-RSP475, from soil samples collected in Talakona, Tirupati, India. Preliminary investigations of its crude extract revealed potent antitumor and xanthine oxidase inhibitory activity, prompting further genomic exploration. To elucidate its biosynthetic potential, we performed whole-genome sequencing, taxonomic classification using ribosomal MLST, and phylogenomic analysis. Additionally, genome annotation was conducted using RAST, TYGS, and BV-BRC platforms. BGC prediction was carried out using antiSMASH, leading to the identification of putative secondary metabolite clusters, including novel and uncharacterized BGCs. These findings underscore the genomic and biosynthetic potential of *Streptomyces* sp. IICT-RSP475 and lay the foundation for future investigations into its novel bioactive metabolites.

## Methods

### Isolation and identification of organism

The strain IICT-RSP475 was isolated from Talakona’s rhizosphere soil in Tirupati, India, using the laboratory-established *i*-Chip method (Tejaswi et al.), and a potent bioactive metabolite with XO inhibition and anti-tumor activities was extracted and purified from culture broth.

### DNA isolation, sequencing and functional genome analysis of IICT-RSP475

DNA was extracted from the pure culture using a HiPurA Bacterial Genomic DNA Purification kit (MB505-250PR HiMedia Laboratories PVT Ltd, Nashik, India) according to the manufacturer’s instructions.

### Whole genome sequencing

Genomic DNA was then sequenced on the Illumina HiSeq 2500 platform. Thus, isolated DNA quality was accessed using the ’Agilent Tape Station’ and the ’Qubit Fluorometer’. Following that, the required amount of DNA was taken as an input and the library was prepared using Illumina’s Nextera DNA Library Prep Kit. The ’ILLUMINA NOVASEQ 6000’ sequencer was used to sequence NGS libraries twice, resulting in about 34 million paired end reads with a read length of 151nt.

### Data analysis

Raw Data Processing and Taxonomic Classification

The raw sequencing data was processed through a series of quality control and filtering steps to ensure high-quality reads for downstream analysis.

#### 1. Quality Assessment and Adapter Trimming

The raw data quality was assessed using FastQC v0.11.7. An adapter contamination range of 3–7% was detected. Nextera transposase adapter sequences (Forward: CTGTCTCTTATACACATCT, Reverse: CTGTCTCTTATACACATCT; Source: ILLUMINA adapters) were removed using Cutadapt v2.8. Reads with a length of ≥50 nucleotides (nt) were retained (Cutadapt parameter: ‘-m 50’). After adapter trimming, the total number of retained paired-end (PE) reads was ∼67 million (67M PE).

#### 2. Taxonomic Classification

The trimmed reads were classified into various taxonomic groups by aligning them to the ‘PlusPFP’ database (maintained by Ben Langmead’s lab) using Kraken v2.0.8. The PlusPFP database comprises all known genomes from bacteria, archaea, viruses, protozoa, fungi, plants, and the human genome. The latest version used in this study was last updated on September 8, 2022. Kraken2 classification results were compiled using in-house scripts and visualized using Krona Tools v2.7.1 (Wood et al. 2019).

These preprocessing steps ensured high-quality, contamination-free data for subsequent genomic analyses.

### Genomic analysis

Primary draft genome analysis was performed by rapid annotation using subsystems technology (RAST) (Brettin et al. 2015) and Bacterial and Viral Bioinformatics Resource Center (BV-BRC) (v3.30.19) platform (Olson et al. 2023).

### Species identification

The ribosomal multi local sequence typing (rMLST) was performed for a precise identification of strain using internal fragments of multiple house keeping genes.

### 16s rRNA sequencing

The molecular characterization of IICT-RSP475 was carries out at Bioserve Biotechnologies Pvt Ltd, Hyderabad, India using 16s rRNA ribotyping. BLASTN version 2.12.0 was used to compare the nucleotide sequence to the NCBI Database. To create the genome BLAST Distance Phylogeny (GBDP) tree, the entire genome sequence was used. Using the OrthoANIu technique, the Average Nucleotide Identity (ANI) of related species with whole genomes was also determined (Yoon et al. 2017).

### Functional Genomic Analysis

Further, the *Streptomyces* sp. IICT-RSP475 draft genome was analysed for secondary metabolite biosynthetic gene clusters using antiSMASH version 5.0 (Blin et al. 2019)

## Results and Discussion

The Kraken2 output for all samples was compiled using in-house scripts. The classification results revealed that 91% of the reads were assigned to bacterial taxa, while 9% remained unclassified (Fig. 1). Using the PlusPFP database, taxonomic analysis identified a single bacterial species in the sequenced sample, classified as an uncharacterized species of *Streptomyces* BH-MK-02 with 76% similarity. To minimize potential false positives, a species richness cutoff of >1% was applied, ensuring that only confidently assigned taxa were considered for further analysis.

**Fig. 1:**
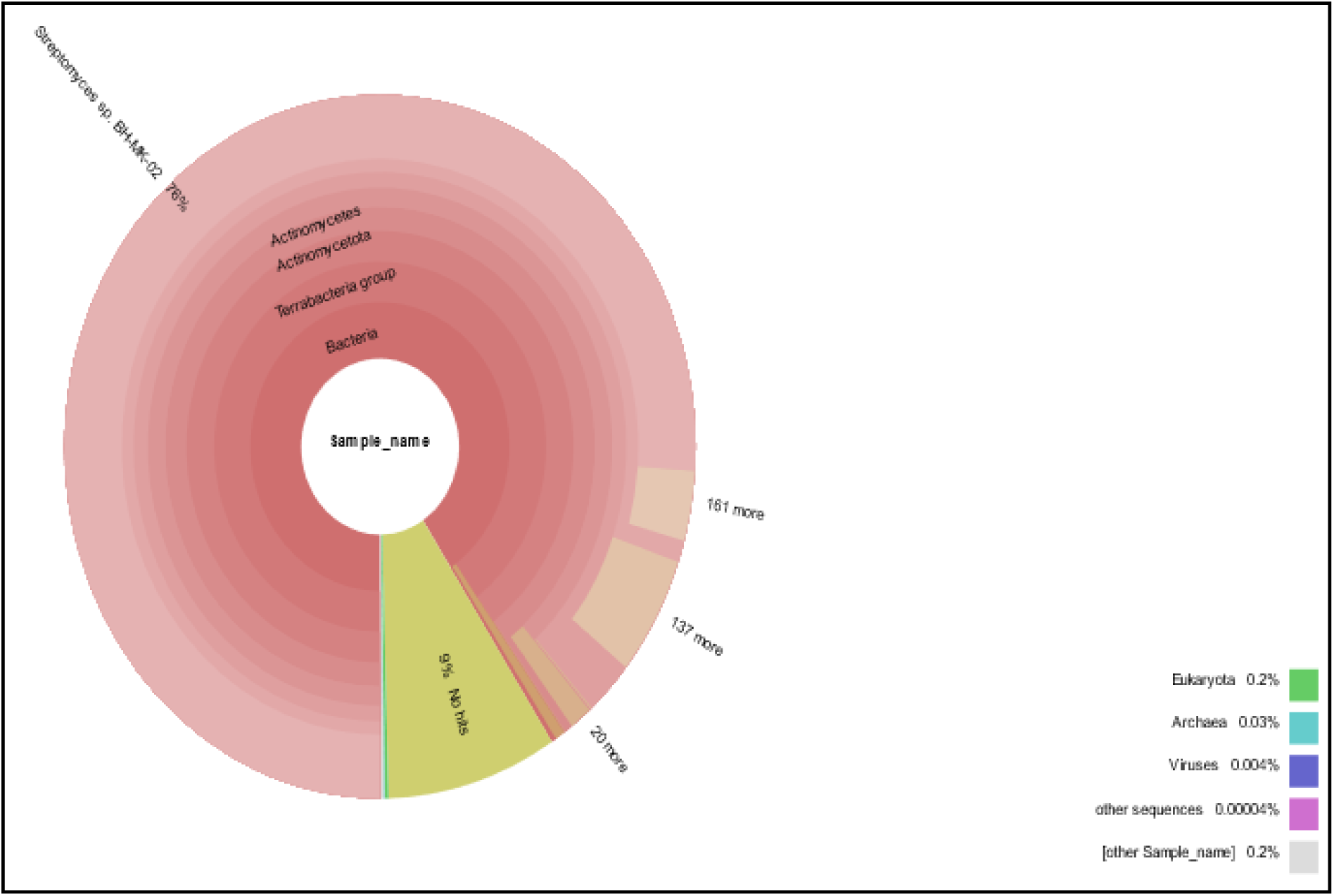
Krona graph representing the DNA sequence reads of *Streptomyces sp.* IICT-RSP475.

### Functional genomic analysis

Primary draft genome analysis was performed by RAST (Brettin et al. 2015) and BV-BRC explored that, there were 750 number of contigs with a total contig size of 4,550,481 bp and N50 contig number of 8,999. The L50 value was 101 and the high GC content of 70.1%. Based on genome annotation, there were 750 number of contigs with protein encoding genes and 145 number of sub systems with 5105 number of coding sequences (Table 1, Fig. 2). There were 6 RNAs.

**Fig. 2:**
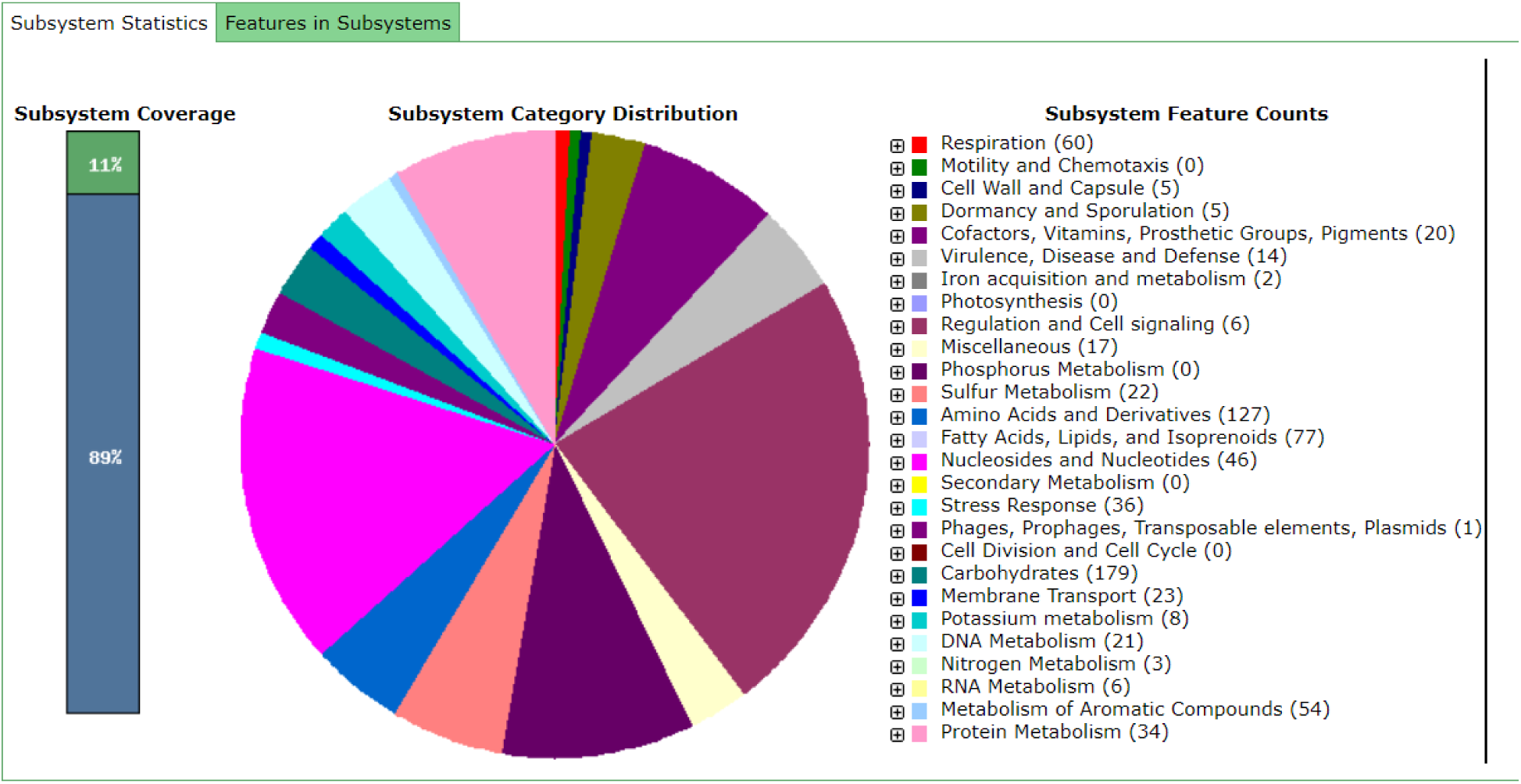
*Streptomyces sp.* IICT-RSP475 subsystem statistics information using RAST annotation. The subsystems category and corresponding feature counts were shown in the legend.

**Table 1:**
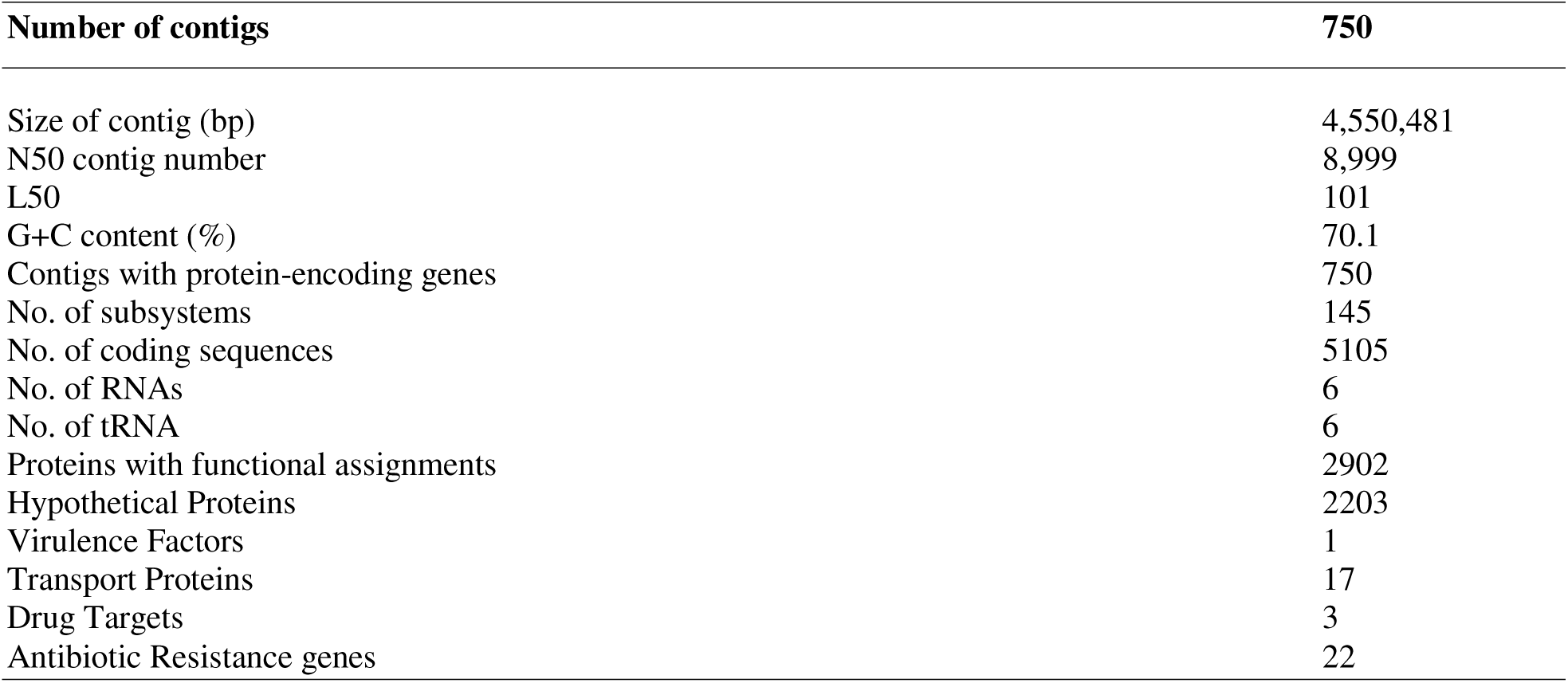
Draft genome characteristics of *Streptomyces sp.* IICT-RSP475 using RAST.

### *Streptomyces sp*. IICT-RSP475 Genome mining for biosynthetic gene clusters (BGCs)

The primary analysis of BGCs from *Streptomyces* sp. IICT-RSP475 genome was analyzed by applying antiSMASH software. This is the most commonly applied bioinformatics tool to find and evaluate BGCs from genome sequences. These studies involves in mining of genome for highlighting the tremendous secondary metabolite BGCS (smBGCs) to discover novel and potentially relevant compounds from the new microbes (Pan et al. 2017). It is evident from earlier genome mining studies that majority of the *Streptomyces* species carry between 8 to 83 BGCs and most commonly distributed classes were non-ribosomal peptide synthetases (NRPS), type 1 polyketide synthases (T1PKS), terpenes (697), other ketide synthases (KS) and lantipeptides. However, butyrolactone, Type 2 PKS (T2 PKS), bacteriocin, and Type 3 PKS (T3 PKS) are the other common BCGs found in *Streptomyces* sp. (Belknap et al. 2020). Similarly, the genome mining of *Streptomyces* sp. IICT-RSP475 in the present study revealed less number of (n=18) BGCs of 3 NRPS, 1 T2 PKS, 3 T3 PKS, 4 terpenes, 5 Ribosomally synthesized and post-translationally modified peptides (RiPPs), N1-siderophore, 1 Cyclodipeptide synthase (CDPS) as depicted in Table 2.

**Table 2:** *Streptomyces sp.* IICT-RSP475 secondary metabolite biosynthesis gene clusters (smBGCs) identified by anti-SMASH.

| Contig | Type | Position (bp) | Cluster known biosynthetic class | Predicted secondary metabolite | Similarity |
| --- | --- | --- | --- | --- | --- |
| Region 2.1 | NAPAA | 1-23,288 | NRP | $\epsilon$ - Poly-L-lysine | 100% |
| Region 2.2 | CDPS | 56,489-77,205 | Unknown | - | - |
| Region 9.1 | Hydrogen-cyanide | 11,486-24,357 | RiPP | Aborycin | 21% |
| Region 11.1 | Terpene | 5,451-26,500 | Polyketide | Ebelactone | 5% |
| Region 39.1 | NRPS | 1-19,391 | NRP | Marformycin A | 25% |
| Region 45.1 | NRPS-like | 1-17,541 | NRP | Kolossin | 100% |
| Region 57.1 | T3PKS | 1-15,399 | Type-III polyketide | Naringenin | 100% |
| Region 60.1 | T3PKS | 1-14,472 | Type-III polyketide | Violapyrone | 28% |
| Region 79.1 | Terpene | 1-10,662 | Unknown | - | - |
| Region 122.1 | RiPP-like | 1-6,985 | Hexacosalactone | Other | 4% |
| Region 187.1 | T3PKS | 1-5,388 | Type-III polyketide | Alkyl resorcinol | 100% |
| Region 233.1 | RiPP-like | 1-4,516 | Unknown | - | - |
| Region 305.1 | T2PKS | 1-3,762 | Type-II Polyketide | Spore pigment | 50% |
| Region 340.1 | Terpene | 1-3,448 | Hopene | Terpene | 15% |
| Region 485.1 | Terpene | 1-2,697 | Hopene | Terpene | 23% |
| Region 544.1 | NI-Siderophore | 1-2,561 | Unknown | Unknown | - |
| Region 611.1 | RiPP-like | 1-2,314 | Unknown | Unknown | - |
| Region 741.1 | RiPP-like | 1-2,021 | Unknown | Unknown | - |

It has been evident from earlier literature that each *Streptomyces* strain of same species varies in their BGC composition. Out of 18 BGCs from *Streptomyces sp.* IICT-RSP475, 12 BGCs showed similarity with known BGCs from antiSMASH database, while 6 are unknown BGCs and represented as non-homologous clusters and these genes did not match in the MIBiG 3.1 known cluster blast. These findings suggest the uniqueness and discovery of novel bioactive compounds from these clusters of *Streptomyces* sp. IICT-RSP475.

The sequence based analysis of BGCs from *Streptomyces* sp. IICT-RSP475 revealed its strong biosynthetic potential along with probability of discovering novel therapeutic compounds from the isolate. The presence of known BGCs contributes its role in antimicrobial activity by inhibiting bacteria, fungi and viruses, anti-inflammatory and anti-cancer potential. They have been associated with secondary metabolites production of Aborycin (Shao et al. 2019) Marformycins (Liu et al. 2015; Bode et al. 2015), naringenin (Álvarez et al. 2015), alkyl resorcinol (Ohnishi et al. 2008), ebelactone (Hong et al. 2016), Violapyrone (Hou et al. 2018), ε - Poly-L-lysine (Purev et al. 2020), Hexacosalactone (Shi et al. 2021).

Whereas, the 6 unknown clusters were classified under 3 BCG clusters. Among them, three clusters belonged to RiPPS, one cluster exhibited N1-siderophore characteristics, one belongs to CDPS cluster and one was in terpene group. The CDPS cluster was found in the region 2.2 consisted of 3 genes associated with biosynthetic process, whereas, N1-siderophore cluster comprised of 24 genes was present in region 544.1 and the terpene cluster was identified in the region 79.1, However, these clusters showed low sequence similarity with characterized CDPS, N1 siderophores and terpene, hence unable to predict the metabolites. In addition, RiPPS-like gene cluster present in the region 233.1 displayed a similarity score of 71% with RiPP-Lanthipeptide from *S. viridochromogenes DSM 40736* which could represents potentially the presence of similar compounds in the genome of *Streptomyces* sp. IICT-RSP475 (Mohimani et al. 2014). Surprisingly, we explored two unidentified RiPPS-like gene clusters in the regions 611.1, 741.1 and did not find any hit in the MIBiG 3.1 known cluster blast, which represents two unique and novel potent metabolites in the genome region. The structures of unidentified BGCs were displayed in the Fig 3.

**Fig 3:**
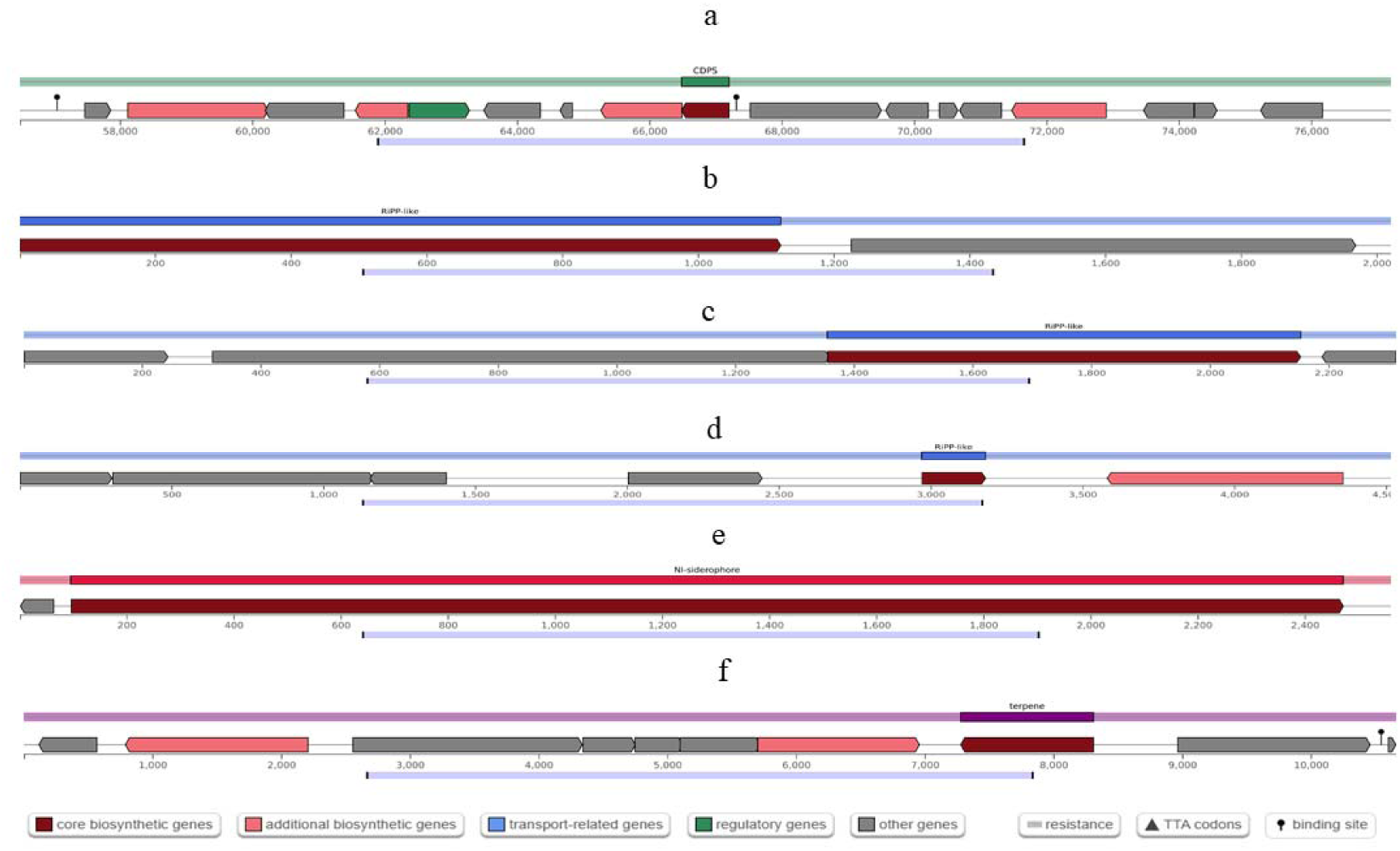
Unidentified biosynthetic gene clusters (BGCs) from Streptomyces sps. IICT-RSP475. a)CDPS, b,c,d) Ripp like, e) N1-siderophore, f) terpene.

Regarding to unknown BGCs, the clusters of CDPS, terpene were associated with gene that synthesizes activators and regulator proteins of antibiotic production. Hence, plays a significant role in the transcription regulation of antibiotic production. These findings explored the novel antibiotic synthesizing gene clusters in the genome of *Streptomyces* sp. IICT RSP-475. Other, unknown clusters need to be discovered by laboratory investigation of metabolites by fermentation.

### 16s rRNA Sequencing

The results from the 16S rRNA sequencing revealed that the strain belongs to the genus *Streptomyces* based on nucleotide homology and phylogenetic analysis. We obtained the 16S rRNA gene sequences of the top 20 nearby species. These results demonstrated that IICT-RSP475 is a member of an unidentified *Streptomyces* species. The Type (Strain) Genome Server (TYGS) independently verified these findings(Meier et al. 2019). The IICT-RSP475 showed 99.73% sequence similarity with unknown species of *Streptomyces* sp. strain IC12A & MM108, as well as with two other species of *Streptomyces* sp. *SXY66* & L4_7_907R. This strain bears close relativeness to *Streptomyces longhuiensis* strain BH-MK-02 and *Streptomyces aureus* strain NBRC 100912. The 16S rRNA gene sequence has been deposited in the NCBI under the Gene accession number OQ612673.

The results of TYGS showed that 16s rRNA gene sequence of IICT-RSP475 strain belongs to the unknown species of *Streptomyces* (Fig 4). Though the strain showed 92.02% ANI with *Streptomyces sporangiiformans* NEAUSSA, it does not considered as *Streptomyces sporangiiformans*. The established cut of values i.e. % dDDH threshold values < 70% and diff G+C content < 1% from TYGS analysis and ANI value < 95-96% indicates that the strain IICTRSP-475 is unique and observed low genomic identity with other species of *Streptomyces* (Table 3). Therefore, it has been observed that the strain warrants further wet lab investigation to identify the expression of novel metabolites.

**Fig 4:**
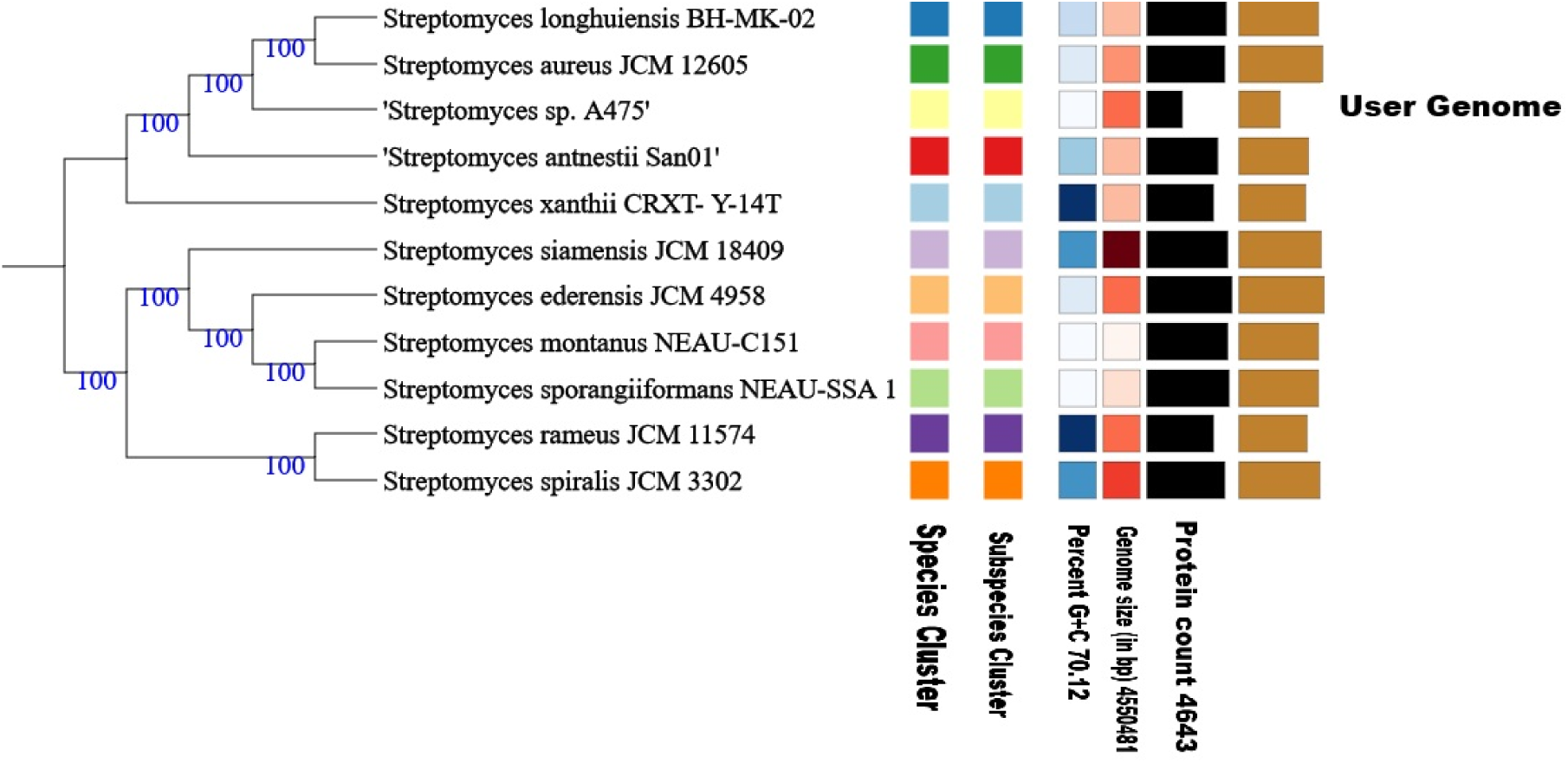
Genome BLAST Distance Phylogeny method (GBDP) diagram of *Streptomyces sp.* IICT-RSP475 generated by using TYGS. Phylogram based on Whole genome sequence.

**Table 3:**
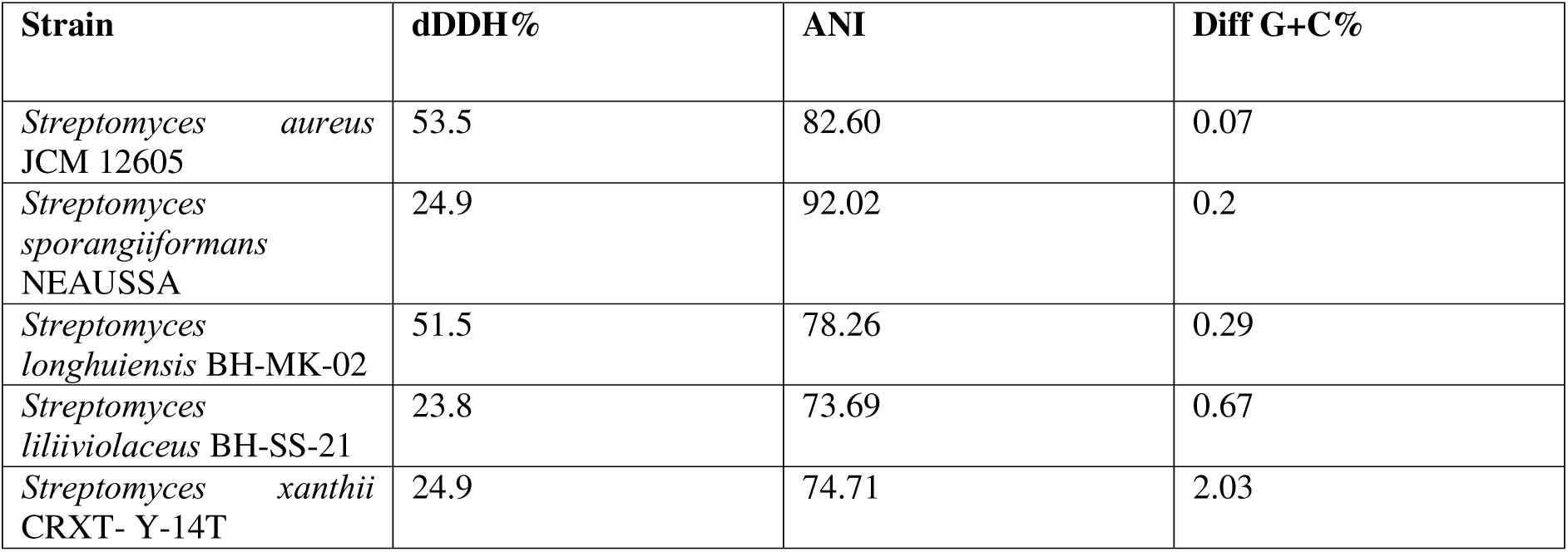
Digital DNA hybridization (dDDH), ANI values and % G+C content of *Streptomyces sp.* IICTRSP-475 with its closely related species using TYGS analysis.

## Data availability

This Whole Genome Shotgun project has been deposited at DDBJ/ENA/GenBank under the accession JBBMGK000000000. The version described in this paper is version JBBMGK010000000.

## Acknowledgments

URB and PRS thank Council for Scientific and Industrial Research (CSIR), New Delhi for funding under the Emeritus Scheme, grant number 21(1102)/20/EMR-II and CSIR-RA fellowship, respectively. Authors thanks to Dr. Linga Banoth, and Ms Parinitha Mandhyani for their constant support. The communication number issued by CSIR-IICT for this article is **IICT/Pubs./2025/092.**

## References

Abbasi MN, Fu J, Bian X, Wang H, Zhang Y, Li A. Recombineering for genetic engineering of natural product biosynthetic pathways. Trends in biotechnology. 2020 Jul 1;38(7):715–28.

Álvarez-Álvarez R, Botas A, Albillos SM, Rumbero A, Martín JF, Liras P. Molecular genetics of naringenin biosynthesis, a typical plant secondary metabolite produced by Streptomyces clavuligerus. Microbial cell factories. 2015 Dec;14:1–2.

Belknap KC, Park CJ, Barth BM, Andam CP. Genome mining of biosynthetic and chemotherapeutic gene clusters in Streptomyces bacteria. Scientific reports. 2020 Feb 6;10(1):2003.

Blin K, Shaw S, Steinke K, Villebro R, Ziemert N, Lee SY, Medema MH, Weber T (2019) antiSMASH 5.0: updates to the secondary metabolite genome mining pipeline, Nucleic Acids Research. 47:W81–W87. 10.1093/nar/gkz310.

Bode HB, Brachmann AO, Jadhav KB, Seyfarth L, Dauth C, Fuchs SW, Kaiser M, Waterfield NR, Sack H, Heinemann SH, Arndt HD. Structure Elucidation and Activity of Kolossin A, the DD/LDPentadecapeptide Product of a Giant Nonribosomal Peptide Synthetase. Angewandte Chemie International Edition. 2015 Aug 24;54(35):10352–5.

Brettin T, Davis JJ, Disz T, Edwards RA, Gerdes S, Olsen GJ, Olson R, Overbeek R, Parrello B, Pusch GD, Shukla M. RASTtk: a modular and extensible implementation of the RAST algorithm for building custom annotation pipelines and annotating batches of genomes. Scientific reports. 2015 Feb 10;5(1):1–6.

Challis, G. L. 2014 Exploitation of the Streptomyces coelicolor A3 (2) genome sequence for discovery of new natural products and biosynthetic pathways. J. Ind. Microbiol. Biotechnol. 41, 219–232)

Hong H, Sun Y, Zhou Y, Stephens E, Samborskyy M, Leadlay PF. Evidence for an iterative module in chain elongation on the azalomycin polyketide synthase. Beilstein Journal of Organic Chemistry. 2016 Oct 11;12(1):2164–72.

Hopwood DA. Highlights of Streptomyces genetics. Heredity. 2019 Jul;123(1):23–32.

Hou L, Huang H, Li H, Wang S, Ju J, Li W. Overexpression of a type III PKS gene affording novel violapyrones with enhanced anti-influenza A virus activity. Microbial Cell Factories. 2018 Dec;17:1–1.

Liu J, Wang B, Li H, Xie Y, Li Q, Qin X, Zhang X, Ju J. Biosynthesis of the anti-infective marformycins featuring pre-NRPS assembly line N-formylation and O-methylation and post-assembly line C-hydroxylation chemistries. Organic letters. 2015 Mar 20;17(6):1509–12.

Meier-Kolthoff JP, Göker M (2019) TYGS is an automated high-throughput platform for state-of-the-art genome-based taxonomy. Nature communications 10:2182. 10.1038/s41467-019-10210-3).

Mohimani H, Kersten RD, Liu WT, Wang M, Purvine SO, Wu S, Brewer HM, Pasa-Tolic L, Bandeira N, Moore BS, Pevzner PA. Automated genome mining of ribosomal peptide natural products. ACS chemical biology. 2014 Jul 18;9(7):1545–51).

Niu G. Genomics-driven natural product discovery in actinomycetes. Trends in biotechnology. 2018 Mar 1;36(3):238–41.

Ohnishi Y, Ishikawa J, Hara H, Suzuki H, Ikenoya M, Ikeda H, Yamashita A, Hattori M, Horinouchi S. Genome sequence of the streptomycin-producing microorganism Streptomyces griseus IFO 13350. Journal of bacteriology. 2008 Jun 1;190(11):4050–60.

Olson RD, Assaf R, Brettin T, Conrad N, Cucinell C, Davis JJ, Dempsey DM, Dickerman A, Dietrich EM, Kenyon RW, Kuscuoglu M. Introducing the bacterial and viral bioinformatics resource center (BV-BRC): a resource combining PATRIC, IRD and ViPR. Nucleic acids research. 2023 Jan 6;51(D1):D678–89.

Pan G, Xu Z, Guo Z, Hindra, Ma M, Yang D, Zhou H, Gansemans Y, Zhu X, Huang Y, Zhao LX. Discovery of the leinamycin family of natural products by mining actinobacterial genomes. Proceedings of the National Academy of Sciences. 2017 Dec 26;114(52):E11131–40.

Purev E, Kondo T, Takemoto D, Niones JT, Ojika M. Identification of ε-poly-l-lysine as an antimicrobial product from an Epichloë endophyte and isolation of fungal ε-PL synthetase gene. Molecules. 2020 Feb 25;25(5):1032.

Shao M, Ma J, Li Q, Ju J. Identification of the anti-infective aborycin biosynthetic gene cluster from deep-sea-derived Streptomyces sp. SCSIO ZS0098 enables production in a heterologous host. Marine drugs. 2019 Feb 21;17(2):127.

Shi P, Li Y, Zhu J, Shen Y, Wang H. Targeted discovery of the polyene macrolide hexacosalactone A from Streptomyces by reporter-guided selection of fermentation media. Journal of Natural Products. 2021 Jun 25;84(7):1924–9.

Tejaswi S, Reddy Shetty P, Siva B, Batchu UR, Chandrasekar C, Reddy PR, Sravanthi V, Banoth L, Suresh Babu K, HMS SK, Jain N. UPLC-Q-TOf-MS Guided Identification of Indole Alkaloids from S. Netropsis IICTRSP-174 Isolated from Leh-Ladakh Soils and Bioactive Profile of Pimprinethine.

Ward AC, Allenby NE. Genome mining for the search and discovery of bioactive compounds: the Streptomyces paradigm. FEMS microbiology letters. 2018 Dec;365(24):fny240.

Wood DE, Lu J, Langmead B (2019) Improved metagenomic analysis with Kraken 2. Genome Biology 20:257.

Yoon SH, Ha S, Lim J, Kwon S, Chun J (2017) A large-scale evaluation of algorithms to calculate average nucleotide identity, Antonie Van Leeuwenhoek, 110: 1281–1286, doi: 10.1007/s10482-017-0844-4.

Zhao Q, Wang L, Luo Y. Recent advances in natural products exploitation in Streptomyces via synthetic biology. Engineering in Life Sciences. 2019 Jun;19(6):452-62.

